## Supplement for "NEMO reshapes the protein aggregate interface and promotes aggrephagy by co-condensation with p62"

### Suppl. Figure 1

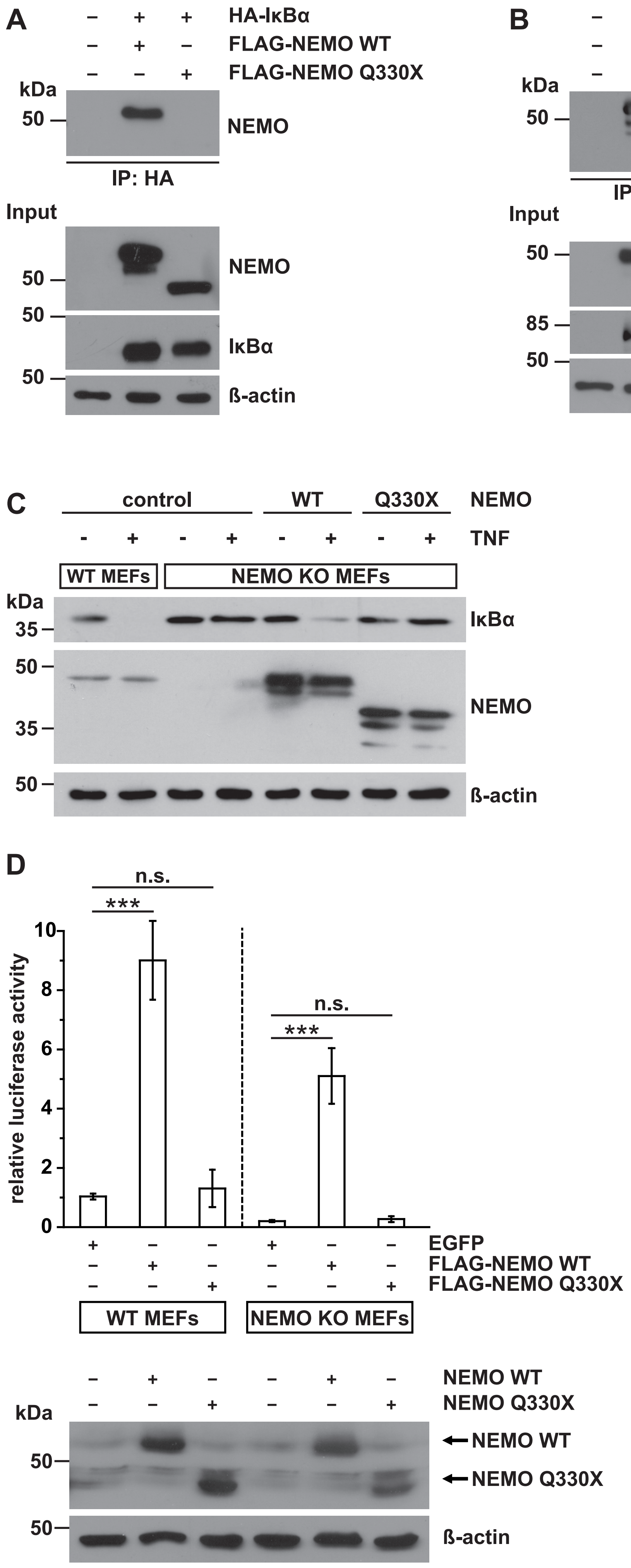

**Figure S1. The Q330X NEMO mutant is defective in NF-κB signaling.**

**A, B. The Q330X mutation disrupts binding of NEMO to IκBα but not to IKKβ.** HEK293T cells were transiently transfected with wildtype FLAG-NEMO or Q330X FLAG-NEMO and HA-IκBα (A) or HA-IKKβ (B) as indicated. One day after transfection, the cells were lysed and HA-tagged proteins were immunoprecipitated using anti-HA-beads followed by immunoblotting using antibodies against NEMO. The input was immunoblotted for NEMO, IκBα (A) or IKKβ (B) and β-actin.

**C. WT NEMO but not Q330X NEMO rescues defective IκBα degradation in NEMO KO MEFs.** WT and NEMO KO MEFs were transiently transfected with WT FLAG-NEMO, Q330X FLAG-NEMO or luciferase as a control. One day after transfection, the cells were treated with TNF (25 ng/ml, 15min) as indicated or left untreated and analyzed by immunoblotting using antibodies against IκBα, NEMO and β-actin.

**D. In contrast to WT NEMO, Q330X NEMO does not promote NF-κB transcriptional activity.** WT and NEMO KO MEFs were transiently transfected with an NF-κB luciferase reporter plasmid and WT FLAG-NEMO, Q330X FLAG-NEMO, or EGFP as a control. 24 h after transfection, the cells were lysed and luciferase activity was measured luminometrically using a plate reader. Data represent the means ± SEM of three independent experiments consisting of three technical replicates each. Statistics: Student’s t-test. Lower panel: Cell lysates were immunoblotted using antibodies against NEMO and β-actin (input control).

Suppl. Figure 2

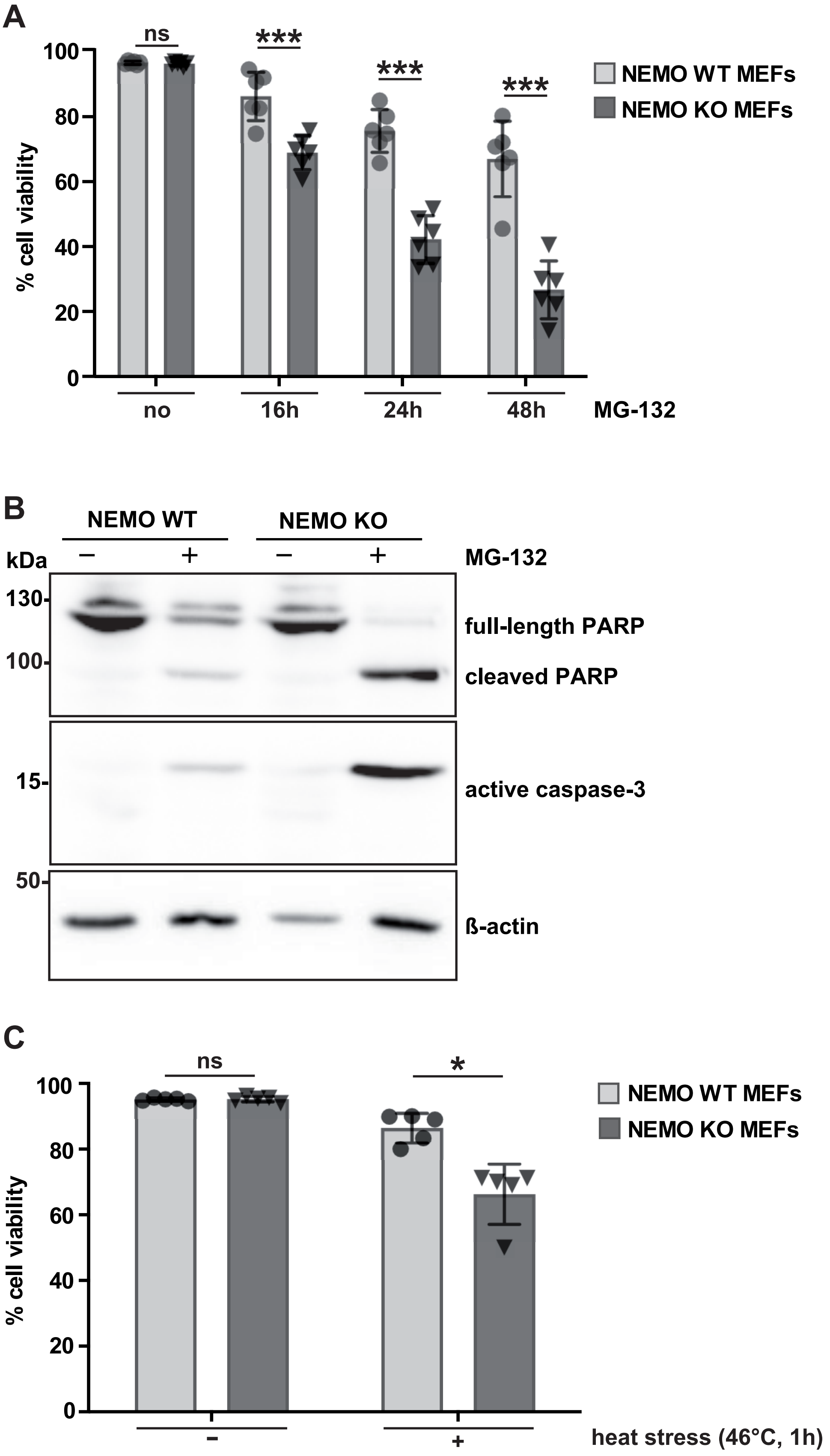

**Figure S2. NEMO KO MEFs are more vulnerable to proteotoxic stress.**

**A. NEMO deficiency decreases cell viability upon proteasomal inhibition.** WT and NEMO KO MEFS were treated with MG-132 for 16 h (2 µM), 24 h (2 µM), or 48 h (0.5 µM), or kept untreated. Cell viability was quantified using Trypan blue dye exclusion. Data are displayed as mean ± SD and were analyzed by two-way ANOVA followed by Bonferroni´s multiple comparison test, n=6.

**B. NEMO deficiency increases apoptotic cell death upon proteasomal inhibition.** WT and NEMO KO MEFS were treated with MG-132 (2 µM, 24 h) and then analyzed by immunoblotting using antibodies against PARP and active caspase-3. The input was immunoblotted for β-actin.

**C. NEMO deficiency decreases cell viability upon heat stress.** WT and NEMO KO MEFS were subjected to a heat stress (46°C, 1 h), followed by overnight recovery. Cell viability was quantified using Trypan blue dye exclusion. Data are displayed as mean ± SD and were analyzed by two-way ANOVA followed by Bonferroni´s multiple comparison test, n=5.

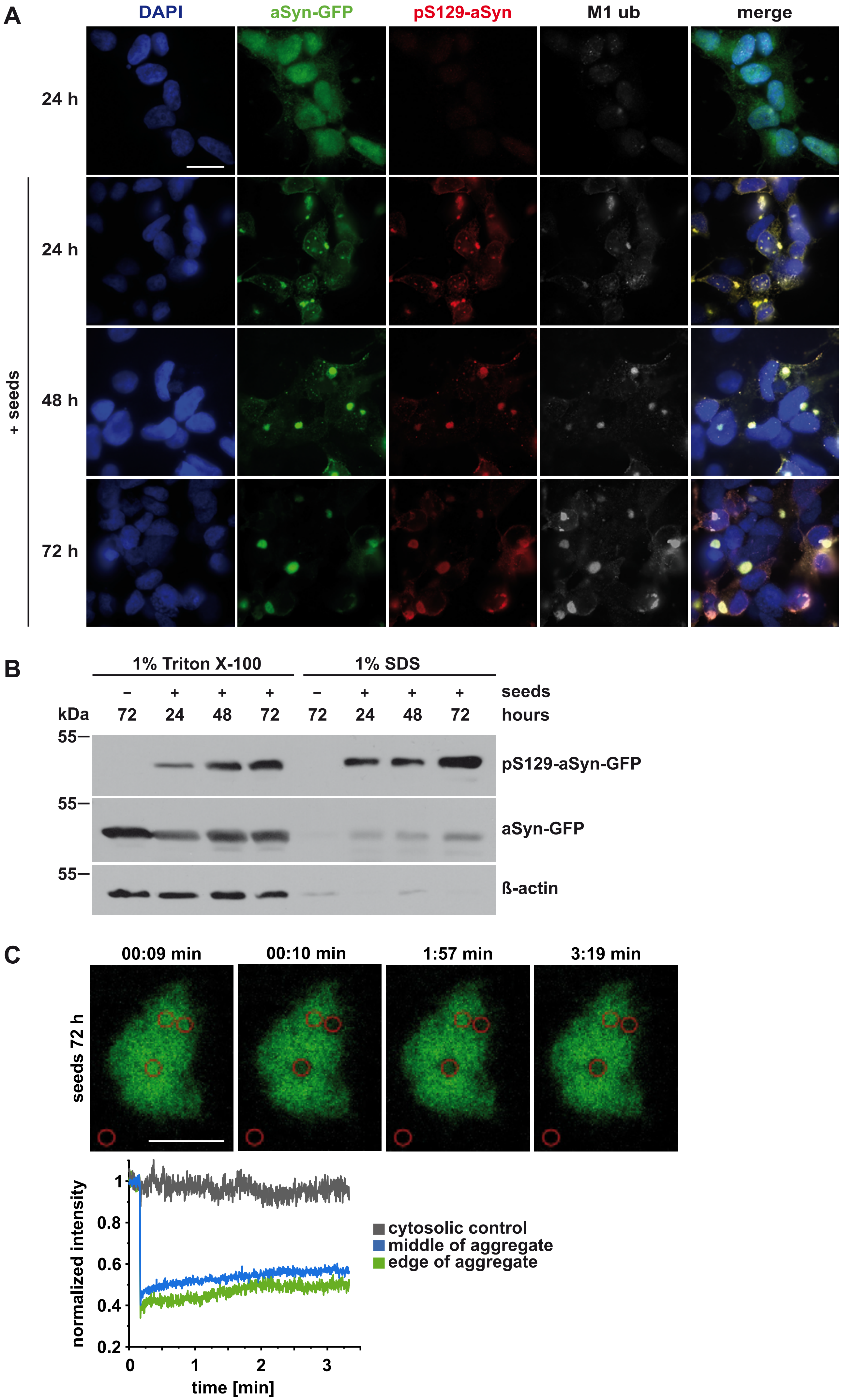
A
DAPI
aSyn-GFP
pS129-aSyn
M1 ub
merge
24 h
24 h
48 h
72 h
+ seeds
B
1% Triton X-100
1% SDS
– + + + – + + +
seeds
kDa
72 24 48 72 72 24 48 72
hours
55—
pS129-aSyn-GFP
55—
aSyn-GFP
55—
ß-actin
C
00:09 min
00:10 min
1:57 min
3:19 min
seeds 72 h
normalized intensity
1
0.8
0.6
0.4
0.2
0
1
2
3
time [min]
cytosolic control
middle of aggregate
edge of aggregate

**Figure S3. aSyn A53T seeds induce aggregation of pS129-positive aSyn-GFP aggregates that colocalize with M1-linked ubiquitin.**

**A. aSyn-GFP aggregates are phosphorylated at S129 and are modified by M1-linked ubiquitin.** SH-SY5Y cells stably expressing aSyn A53T-GFP were treated with aSyn A53T seeds (+ seeds) or PBS as a control, fixed 24 h, 48 h, or 72 h after seeding, and analyzed by immunocytochemistry and fluorescence SR-SIM using antibodies against pS129-aSyn and M1-linked ubiquitin. Scale bar, 20 µm.

**B. aSyn A53T seeds induce aggregation and phosphorylation of aSyn-GFP at S129.** SH-SY5Y cells stably expressing aSyn A53T-GFP were treated with aSyn A53T seeds or PBS as a control and harvested 24 h, 48 h, or 72 h after seeding. The cells were subjected to a detergent solubility assay and analyzed by immunoblotting using antibodies against pS129-aSyn, aSyn, and β-actin.

**C. aSyn A53T seeds induce the formation of immobile aSyn-GFP aggregates.** SH-SY5Y cells stably expressing aSyn A53T-GFP were treated with aSyn A53T seeds and GFP aggregates in living cells were analyzed by FRAP (fluorescence recovery after photobleaching) 72 h after seeding. An area in the middle and at the edge of an aSyn aggregate was photobleached. Representative images are shown in the upper panel. Scale bar, 5 µm. Normalized intensities of the FRAP experiments are shown in the lower panel. All data are displayed as mean, n=3.

### Suppl. Figure 4

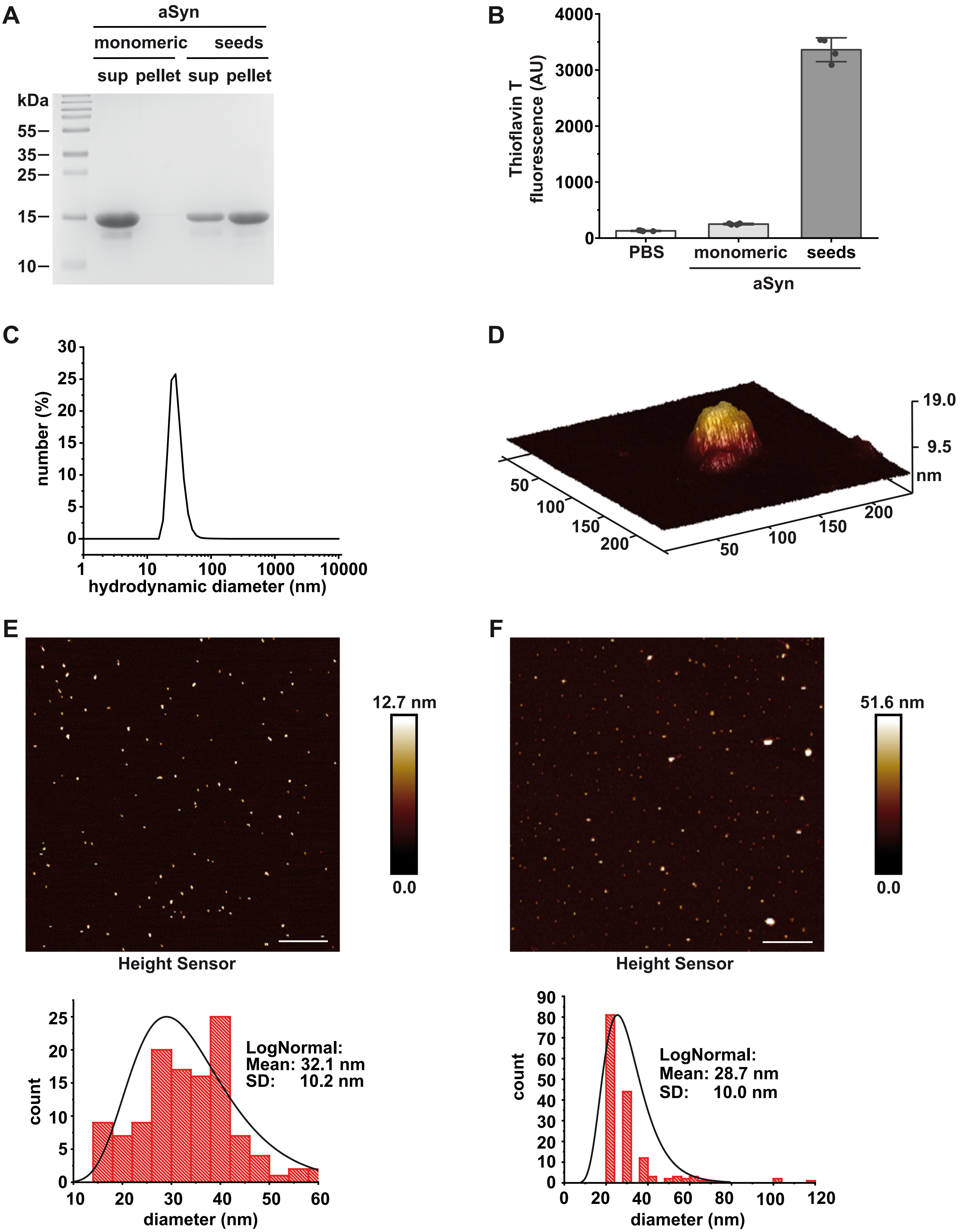

**Figure S4. Validation of aSyn A53T seeds.**

**A. Sedimentation assay for monomeric aSyn A53T and aSyn A53T seeds.** *In vitro* formed aSyn A53T seeds were separated from soluble aSyn A53T monomers by centrifugation. Supernatant (sup) and pellet fractions were analyzed by SDS-PAGE and Coomassie staining.

**B. aSyn A53T seeds bind to Thioflavin T.** Fluorometry of monomeric aSyn, aSyn seed or PBS as a control was performed using a microplate reader. Data are shown as mean ± SD, n=4 technical replicates for each sample.

**C. Derived hydrodynamic diameter of aSyn A53T seeds measured by dynamic light scattering (DLS).** The size distribution of freshly sonicated aSyn A53T seeds dispersed in PBS was evaluated from the derived intensity, considering the number distribution of colloids. A mean diameter of 27.7 nm with a standard deviation of 5.7 nm was derived from a Gaussian fit.

**D. Analysis of aSyn A53T seeds by atomic force microscopy (AFM).** 3D rendered image of a single aSyn A53T seed adsorbed on MICA from a freshly sonicated dispersion of aSyn A53T seeds in PBS, measured by liquid AFM in peak force mode. Rendering was done on a 240x240 nm height sensor image by NanoScope Analysis software from Bruker. The measured feature exhibits a vertical length of 59 nm, a horizontal length of 41 nm and a maximum height of 10 nm.

**E. aSyn size distribution of LAFM height measurements on MICA in PBS solution.** Obtained 2D mapping of the measured surface (Top: size 6.6 x 6.6 µm). Particle size distribution was derived by particle analysis mode of the Bruker Nanoscope software: Height threshold was set to > 5 nm. Statistical distribution was evaluated with a Log normal function: Mean particle diameter 32.1 ± 10.2 nm.

**F. aSyn size distribution of AFM height measurements on Si-wafer after 4 µl dropcasting.** Obtained 2D mapping of the measured surface (Top: size 6.6 x 6.6 µm). Particle size distribution was derived by particle analysis mode of the Bruker Nanoscope software: Height threshold was set to > 16 nm. Statistical distribution was evaluated with a Lognormal function: Mean particle diameter 28.7 ± 10.0 nm.

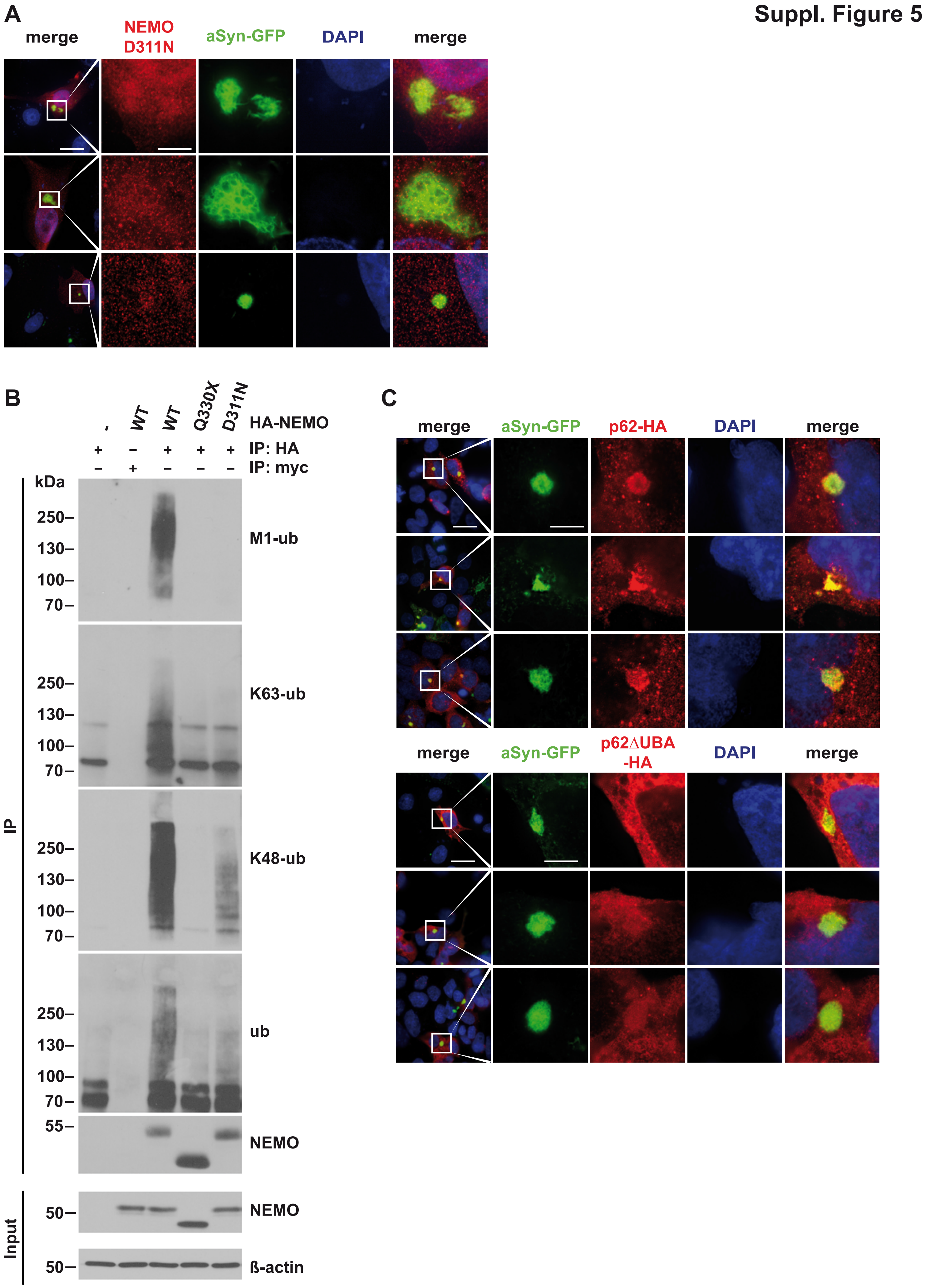
Suppl. Figure 5
A
merge
NEMO D311N
aSyn-GFP
DAPI
merge
B
-
WT
WT
Q330X
D311N
HA-NEMO
+ − + + + IP: HA
− + − − − IP: myc
kDa
250–
130–
100–
70–
M1-ub
K63-ub
K48-ub
ub
55–
NEMO
50–
NEMO
ß-actin
IP
Input
C
merge
aSyn-GFP
p62-HA
DAPI
merge
p62∆UBA -HA

**Figure S5. Impaired recruitment of D311N NEMO and p62ΔUBA to aSyn aggregates.**

**A. D311N NEMO is not recruited to aSyn aggregates.** SH-SY5Y cells stably expressing aSyn A53T-GFP were transiently transfected with D311N HA-NEMO. After 24 h, the cells were treated with aSyn A53T seeds, fixed 48 h after seeding, and analyzed by immunocytochemistry and fluorescence SR-SIM using an antibody against the HA-tag. Scale bar, 20 and 5 µm.

**B. Modification of NEMO Q330X with ubiquitin is impaired**. HEK293T cells were transiently transfected with plasmids encoding HA-tagged wildtype NEMO, Q330X NEMO, or D311N NEMO. Cells were lysed under denaturing conditions and the HA-tag was immunoprecipitated using anti-HA agarose; anti-myc-agarose served as a control for unspecific binding. Samples were analyzed by immunoblotting with antibodies specific for M1-, K63-, K48-linked, or pan-ubiquitin.

**C. Recruitment of p62 lacking the UBA domain aSyn aggregates is impaired**. SH-SY5Y cells stably expressing aSyn A53T-GFP were transiently transfected with plasmids encoding either HA-tagged p62 or p62ΔUBA as indicated. After 24 h, the cells were treated with aSyn A53T seeds, fixed 48 h later, and analyzed by immunocytochemistry and fluorescence SR-SIM using an antibody against the HA-tag. Scale bar, 20 and 5 µm.

| Pathology | Brain region | Identifier | Type | source |
|---|---|---|---|---|
| PD | midbrain | T92/174 | Paraffin | Charité, Berlin, Germany |
| AD | frontal isocortex | A133/17 | Paraffin | Institute of Neuropathology, University Medical Center, Hamburg-Eppendorf, Germany |
| FTLD | frontal isocortex | 199 | Paraffin | |
| DLBD | frontal isocortex | A199/17, A206/16 | Paraffin | |
| NEMO patient | middle frontal gyrus | A16-153 | Paraffin | Department of Pathology, University of California, San Francisco, California, USA. |
| DLBD | middle frontal gyrus | A17-159 | Paraffin | |
| control | middle frontal gyrus | A17-47 | Paraffin | |

**Table S1.** ***Post mortem*** **brain samples data**
